# Compressed sensing-based super-resolution ultrasound imaging for faster acquisition and high quality images

**DOI:** 10.1101/2020.12.18.423443

**Authors:** Jihun Kim, Qingfei Wang, Siyuan Zhang, Sangpil Yoon

**Affiliations:** Department of Aerospace & Mechanical Engineering, University of Notre Dame, Notre Dame, IN, USA; Department of Biological Sciences, University of Notre Dame, Notre Dame, IN, USA

**Keywords:** Super-resolution ultrasound imaging, Compressed sensing, Angiogenesis, microvessel imaging

## Abstract

Super-resolution ultrasound (SRUS) imaging technique has overcome the diffraction limit of conventional ultrasound imaging, resulting in an improved spatial resolution while preserving imaging depth. Typical SRUS images are reconstructed by localizing ultrasound microbubbles (MBs) injected in a vessel using normalized 2-dimensional cross-correlation (2DCC) between MBs signals and the point spread function of the system. However, current techniques require isolated MBs in a confined area due to inaccurate localization of densely populated MBs. To overcome this limitation, we developed the ℓ_1_-homotopy based compressed sensing (L1H-CS) based SRUS imaging technique which localizes densely populated MBs to visualize microvasculature in vivo. To evaluate the performance of L1H-CS, we compared the performance of 2DCC, interior-point method based compressed sensing (CVX-CS), and L1H-CS algorithms. Localization efficiency was compared using axially and laterally aligned point targets (PTs) with known distances and randomly distributed PTs generated by simulation. We developed post-processing techniques including clutter reduction, noise equalization, motion compensation, and spatiotemporal noise filtering for in vivo imaging. We then validated the capabilities of L1H-CS based SRUS imaging technique with high-density MBs in a mouse tumor model, kidney, and zebrafish dorsal trunk, and brain. Compared to 2DCC, and CVX-CS algorithm, L1H-CS algorithm, considerable improvement in SRUS image quality and data acquisition time was achieved. These results demonstrate that the L1H-CS based SRUS imaging technique has the potential to examine the microvasculature with reduced acquisition and reconstruction time of SRUS image with enhanced image quality, which may be necessary to translate it into the clinics.

## I. Introduction

CONTRAST enhanced ultrasound imaging has been developed using microbubbles (MBs) to provide improved therapeutic outcomes and anatomical and functional information for the diagnosis of disease and preclinical investigations. MBs have been utilized for blood-brain barrier opening and drug/gene delivery by stable and inertial oscillations to increase the permeability of connective tissues in brain vasculature [1–3]. Targeting specific biomarkers using MBs with functional moieties detects tumor angiogenesis with high specificity using ultrasound imaging [4, 5]. Recently, a series of development of MB mediated ultrasound imaging techniques tries to overcome the diffraction limit of conventional ultrasound imaging while preserving penetration depth with improved detection capability for clinical and preclinical applications. Super-resolution ultrasound (SRUS) imaging technique utilizes the blinking of MBs by compounding thousands of frames to construct microvessel images to examine the structural abnormalities and dysfunction of the vessel in various types of organs and leaky vasculature in tumor mass [6–12]. Before constructing the SRUS image, the clutter signals of each frame are removed by applying to transmit or post-processing techniques such as pulse inversion [13], differential imaging [14], and singular value decomposition (SVD) filtering [15]. Among them, SVD filtering was demonstrated as a suitable processing step to precisely localize MBs in the range of few micrometers [16].

Normalized 2-dimensional cross-correlation (2DCC) technique is then utilized to localize MBs by comparing detected MBs signals and the point-spread function (PSF) of the ultrasound imaging system [7]. However, 2DCC algorithm does not accurately detect densely populated MBs. Reduced MBs concentration may be used to avoid inaccurate localization due to dense MBs, which increase data acquisition time. For example, a total of 75,000 frames at a frame rate of 500 Hz was acquired to reconstruct the SRUS image of rat brain vasculature within 150 sec [17]. The accompanied drawback with longer acquisition time may cause misalignment between frames due to the motion of an imaging target. Recently, an isolation technique using a 3D Fourier transform was introduced to overcome this limitation by dividing the subset of MBs signal according to the flow velocities and directions at the region of interest (ROI) in a vessel [18]. However, it is challenging to determine optimal subsets due to the unpredictable flow rate in deeper microvessel *in vivo*, which may cause blurring and distortion of original MBs signals and inaccurate localization of MBs.

Recently, compressed sensing (CS) has been introduced as a promising technique to reconstruct original signals from sparsely sampled signals that do not satisfy the Nyquist-Shanon sampling theorem [19]. CS has shown great success in accelerating the magnetic resonance imaging by allowing the irregular undersampling [20]. Especially, CS-based stochastic optical reconstruction microscopy (STORM) was utilized for the reconstruction of a super-resolution optical image to visualize microtubule dynamics in living cells. This showed the capability to recover closely located structures labeled with fluorophores [21]. Furthermore, CS algorithm was demonstrated that this could be utilized for the localization of high-density MBs for the reconstruction of SRUS images in simulation and in vivo application while reducing the acquisition time of ultrasound image frames [22, 23]. However, the standard CS algorithm based on the interior point method is computationally expensive, thus, the computational time was measured as ~7.58 hours/frame with an image size of 64 × 64 pixels for SRUS imaging [23]. To solve the minimization problem efficiently for the rapid CS algorithm, alternative methods were introduced [24, 25]. This alternative method based CS algorithm was applied for SRUS imaging of the rabbit kidney in vivo but utilized the constant noise parameter to solve the minimization problem [26]. The ℓ_1_-homotopy based CS (L1H-CS) algorithm showed the capability to search the optimized parameter naturally [27]. Fast super-resolution optical imaging of microtubules in BS-C-1 cells was performed based on the L1H-CS algorithm, which showed a ~300-fold faster computational time than the standard CS algorithm [28].

In this paper, we present an L1H-CS based SRUS imaging technique that could reduce both the computational time and the acquisition time of ultrasound image frames for the reconstruction of the SRUS image. The localization efficiency, error, and computational time of the proposed technique were evaluated by comparing it with the conventional localization technique using the two and randomly distributed point targets (PTs) generated in simulation. Furthermore, we developed clutter reduction, noise equalization, motdion correction, and spatiotemporal noise filtering algorithms to perform in vivo imaging. The contrast to tissue ratio (CTR) and signal to noise ratio (SNR) were then analyzed quantitatively at ROI. Besides, we performed SRUS imaging of mouse tumor, kidney, and zebrafish dorsal trunk, and brain with the injection of high-density MBs. The number of frames and computational time required for the reconstruction of in vivo microvasculature images based on the proposed and conventional techniques were examined. Results demonstrate that the SRUS imaging technique could be substantially improved by applying L1H-CS algorithm for the localization of high-density MBs, suggesting its potential as a faster SRUS imaging technique.

## II. Materials and methods

### A. ℓ_1_-homotopy based compressed sensing

Mathematically, detected MBs signals, *y*, using the ultrasound imaging system have a linear relationship with MBs location, *x*

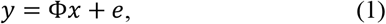

where *y* and *x* consist of the one column vectors reshaped from the 2D matrix by row-wise concatenating the matrix of the original image or the localized results in the upsampled space or pixels [Fig. 1(a)], respectively. The matrix Φ represents the point spread function (PSF) derived by an imaging system by simulation or actual measurements. *e* denotes a noise vector. The ith column of matrix Φ corresponds to the acquired image when only a one-point target exists at a particular index i of *x*. To achieve upsampled localized results, *x*, at a measured frame, *y*, the interior point method utilizes to resolve the ℓ_1_-norm minimization problem for CS as follows [21]:

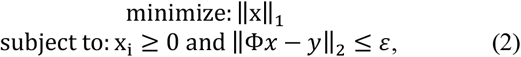

where *ε* denotes a constant showing the balance of fidelity and sparsity in the optimization algorithm. For example, a perfect fit for an original image is estimated when the *ε* value is 1, but *ε* is generally set somewhat larger than 1 to apply uncertainty to the variance measurement. For in vivo imaging we selected *ε* to be 2.3 [21]. In this paper, the interior point method based CS algorithm was implemented by using the CVX optimization package in Matlab (CVX-CS) [29].

**Fig. 1.**
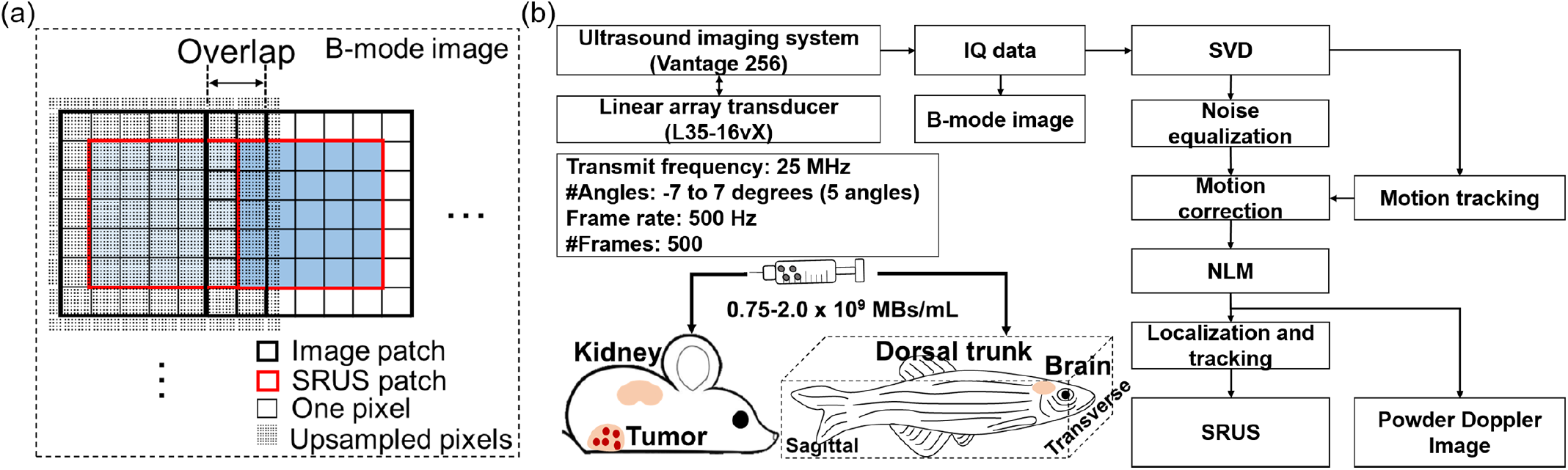
(a) For compressed sensing-based reconstruction of SRUS image, upsampled image patches (thick black square) were used by overlapping 2 pixels between adjacent image patches to remove false localization of point targets near boundaries to correctly build SRUS patches (red thick square). By stitching SRUS patches side by side, a complete SRUS image was acquired from B-mode image (black dashed square). (b) The schematic picture describes overall post-processing steps to reconstruct SRUS images and the administration of microbubbles to mouse tumor models and zebrafish.

To solve the minimization problem efficiently, an alternative method was proposed as follows [24, 25]:

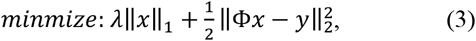

where *λ* is a positive weight. Eq. (3) was expanded to build the L1H-CS formulation using the homotopy parameter, *ϵ*, as follows [27, 30]:

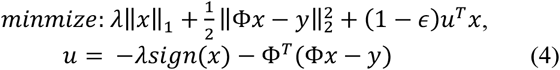

As *ϵ* adjusts from 0 to1, the Eq. (4) steadily transforms to the Eq. (3). The homotopy method offers a typical framework that can solve an optimization problem by continuously transforming it into a related problem with less computing cost [27]. For the reconstruction of SRUS image based on L1H-CS. In this paper, we utilized the L1H-CS package provided in https://github.com/sasif/L1-homotopy [27].

To reduce the computing cost of the CVX-CS, and L1H-CS algorithms, image patches with a kernel size of 7 × 7 pixels were used [thick black square in Fig. 1(a)]. Each image patch was extended by half pixels after upsampled by the factor of 8. Adjacent image patches were overlapped by 2 pixels to localize MBs by CS algorithms. Upsampled image patches were stitched patch by patch after taking 5 × 5 pixels to form SRUS patch [thick red square in Fig. 1(a)] to prevent false localization of MBs close to edges of image patches. The image patch-based CS algorithms allowed parallel computing in Matlab to improve computing speed substantially. The computation is performed on a personal desktop, with an AMD Ryzen 7 3700X 3.6 GHz processor and 16 GB RAM.

### B. Ultrasound imaging configuration

For the acquisition of the in-phase and quadrature (IQ) data, we utilized an ultrasound imaging research platform (Vantage 256, Verasonics Inc., Kirkland, WA, USA) with a high-frequency linear array transducer (L35-16vX, Verasonics Inc., Kirkland, WA, USA). The transducer was incorporated with a 3-axis motorized stage (ILS150CC, Newport Corp., Irvine, CA, USA). A five angle (from −7° to 7° with a step size of 3.5°) plane-wave imaging with an effective frame rate of 500 Hz and transmit frequency of 25 MHz was used. The excitation peak-to-peak voltage of 10 V with a one-cycle sinusoidal signal was generated for each transmission angle. A total of 500 compounded IQ data sets were acquired for 1 sec with pixel resolutions of 27.5 and 34.5 μm in axial and lateral directions, respectively. Furthermore, PSF at the elevation focus of ~ 8 mm was acquired using the Verasonics simulator. The PSF acquired in the simulation was utilized to evaluate localization techniques using two-PTs and randomly distributed PTs. For in vivo imaging, the PSF of MBs was generated as follows:

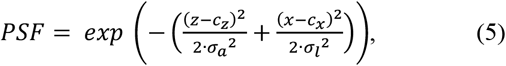

where (*Z*, *x*) and (*C*_z_, *C*_x_) consist of the coordinate and center position of the generated PSF. The σ_a_ and σ_l_ represent the average full width at half maximum (FWHM) of MBs in axial and lateral directions, respectively. In this work, the average FWHM for each direction was measured by manually selecting 10 MBs at different frames. The average FWHM of PSF in axial and lateral direction was measured to be 39.6 and 48.7 μm, respectively.

### C. Post-processing procedures for in vivo super-resolution ultrasound imaging

The post-processing procedures for SRUS imaging in vivo are shown in Fig. 1(b). To remove the clutter signal, SVD based spatiotemporal filtering was applied to the IQ data. The SVD algorithm was briefly summarized as follows. First, the stacked spatial IQ data are converted into a 2D space-time matrix called Casorati matrix *S*. The SVD of *S* was described as follows [15]:

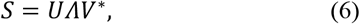

where the column of *U* and *V* matrices are the associated spatial singular and temporal vectors, and *\** represents the conjugation transpose. *Λ* is a diagonal matrix that consists of singular values. In this paper, the SVD filtering was implemented using the “*svd.m*” in Matlab. To remove the clutter signals, the lower (*T_l_*) value thresholds were set by the examination of the turning point on the singular value orders [31, 32]. Then, the filtered signal was reconstructed by:

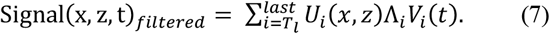

After removing the clutter signals, IQ data sets were converted to the intensity data by the square root of the sum of squares. Then, the noise equalization algorithm was applied to compensate for the ramp-shaped background noise which might occur due to the varying gain along with the depth [33]. The background noise was derived by the reconstructed signal using only the lowest singular value. Then, the noise equalized MBs images were extracted by dividing the background noise profile. The rigid motion of noise equalized MBs images were corrected. The rigid motion in axial and lateral direction throughout the frames at the ROI was estimated by applying the subwavelength motion correction algorithm with the first frame of clutter signals as a reference [34]. To further reduce the noise while preserving the MBs signal, we applied nonlocal means (NLM) filter to the spatiotemporal signals [7].

MBs were then localized by applying the CVX-CS, and L1H-CS algorithms to denoised MBs signals as described in the previous section. We also developed 2DCC based localization technique to compare the performance of CS-based localization technique. Denoised MBs and PSF signals were spatially interpolated 8× under the cubic spline interpolation. 2DCC between the interpolated MBs and PSF signals was conducted. The centroid of MBs was found using the regional peak points finding algorithm (“imregionalmax.m” in Matlab). These processes were iterated for every frame. Each MB localized in each frame was tracked by pairing with the nearest MB in the following frames using the Hungarian algorithm (https://github.com/tinevez/simpletracker) [35, 36]. Finally, the tracked MBs were superimposed to reconstruct the final SRUS image. Furthermore, the Power Doppler (PWD) image was produced using the denoised MBs signal as follows [15]:

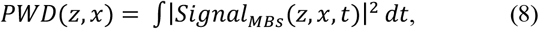

To evaluate denoising techniques including SVD, SVD + noise equalization, SVD + noise equalization + NLM filtering, we measured CTR and SNR of denoised images as follows:

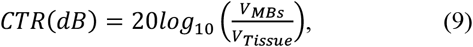

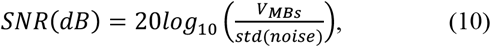

where *V*_*MBS*_ and *V*_*Tissue*_ are the signals of MBs and tissues, respectively. The std(noise) represents the standard deviation of the noise signal. Besides, to compare the localization techniques quantitatively, we defined the vessel density (VD) in the tumor as follows [37].

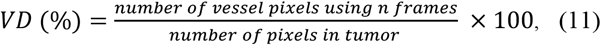

Tumor areas were selected manually by examining the B-mode and SRUS images of mouse tumors.

### D. Point target preparation using simulation

To evaluate the performance of localization using 2DCC, CVX-CS, and L1H-CS algorithms, two points aligned along with axial and lateral directions and randomly distributed PTs were generated in simulation. First, two-PTs were constructed at each inter-point distance from 13.75 to 110 μm with a step size of 13.75 μm in axial and lateral directions, respectively. The wavelength in this study was 61.6 μm. We localized these two points using 2DCC, CVX-CS, and L1H-CS algorithms to examine the minimum distance that can be localized in axial and lateral directions.

To evaluate localization efficiency, error, and computational time, we generated randomly distributed PTs with different densities at 2.65, 7.95, 15.89, 26.48, 39.73, 55.62, 74.16, 95.35, 119.2, 145.7, and 174.8 PTs/mm^2^. Localized PTs were paired with the nearest PTs generated in simulation using Hungarian pairing algorithm with a limitation of a maximum pairing distance of 50 μm. Then, the localization efficiency and errors were identified by counting the localized PTs and calculating the distance deviated from the locations of simulated PTs. These were iterated 5 times for each density. Then, the average localization efficiency and error with the standard deviation of 2DCC, CVX-CS, and L1H-CS algorithms were measured (n = 5). Furthermore, we compared the computational time with an image size of 62 × 62 pixels at each density of PTs.

### E. In vivo imaging of mouse tumor model and normal adult zebrafish

The proposed technique was validated by SRUS imaging of mouse and zebrafish. The animal experimental protocols including mouse and zebrafish were approved by the Institutional Animal Care and Use Committee of the University of Notre Dame.

Mammary tumor model (C57BL/6, Stock No:000664, The Jackson Laboratory, Bar Harbor, ME, USA) was generated by injection of 500K E0771 cells resuspended in PBS with Matrigel (BD Biosciences, San Jose, CA, USA) at 1:1 ratio into the fat pad of the abdominal mammary gland, and imaging was performed 10 to 14 days post-injection. A ketamine/xylazine cocktail was intraperitoneally injected for anesthesia. Hairs from the ROI were removed using the depilatory cream (Nair®). The MBs (Lumason, Bracco Diagnostics Inc., Princeton, NJ, USA) with a volume of 150 μL (0.75 ‒ 2.0 × 109 MBs/mL) were injected through the retro-orbital of the mouse. For the retro-orbital injection of MBs to the mouse, we utilized a 1 mL syringe with a 27 g beveled needle.

We further performed the SRUS imaging of normal adult zebrafish (LiveAquaria, Rhinelander, WI, USA) aged 3 to 12 months. To anesthetize zebrafish, the ethyl 3-aminobenzoate methanesulfonate (MS-222, Sigma-Aldrich, St. Louis, MO, USA) was used. The zebrafish were anesthetized by immersion in 4 % anesthetic. MBs with a volume of 3 μL (0.75 ‒ 2.0 × 109 MBs/mL) was injected intraperitoneally using a 10 μL syringe (NANOFIL, World Precision Instruments, Sarasota, FL, USA) with a 33 g beveled needle. [38].

## III. Results

### A. Evaluation of the localization techniques using two-point targets in simulation

The performance of the localization capability was analyzed by comparing 2DCC, CVX-CS, and L1H-CS based localization techniques. Two-PTs separated by a known distance in axial and lateral directions were used. Figs. 2(a-f) show the minimum distance between two-PTs that can be localized by 2DCC, CVX-CS, and L1H-CS algorithms, respectively. These were overlaid with B-mode images. Blue, green, and red asterisks indicate the locations of localized points by 2DCC, CVX-CS, and L1H-CS, respectively. Black circles indicate the location of simulated points. Figs. 2(g-h) present the number of localized points depending on the distance between two points by 2DCC, CVX-CS, and L1H-CS. Because only two-PTs were used, the maximum number of localizations was 2. 2DCC localizes two simulated points correctly when the distance between two points becomes 82.5 μm in the axial direction and 96.3 μm in the lateral direction. CVX-CS can localize two points when they are 41.3 μm apart from each other in axial and lateral directions. L1H-CS algorithm can localize two points when the separation is 27.5 μm in both directions.

**Fig. 2.**
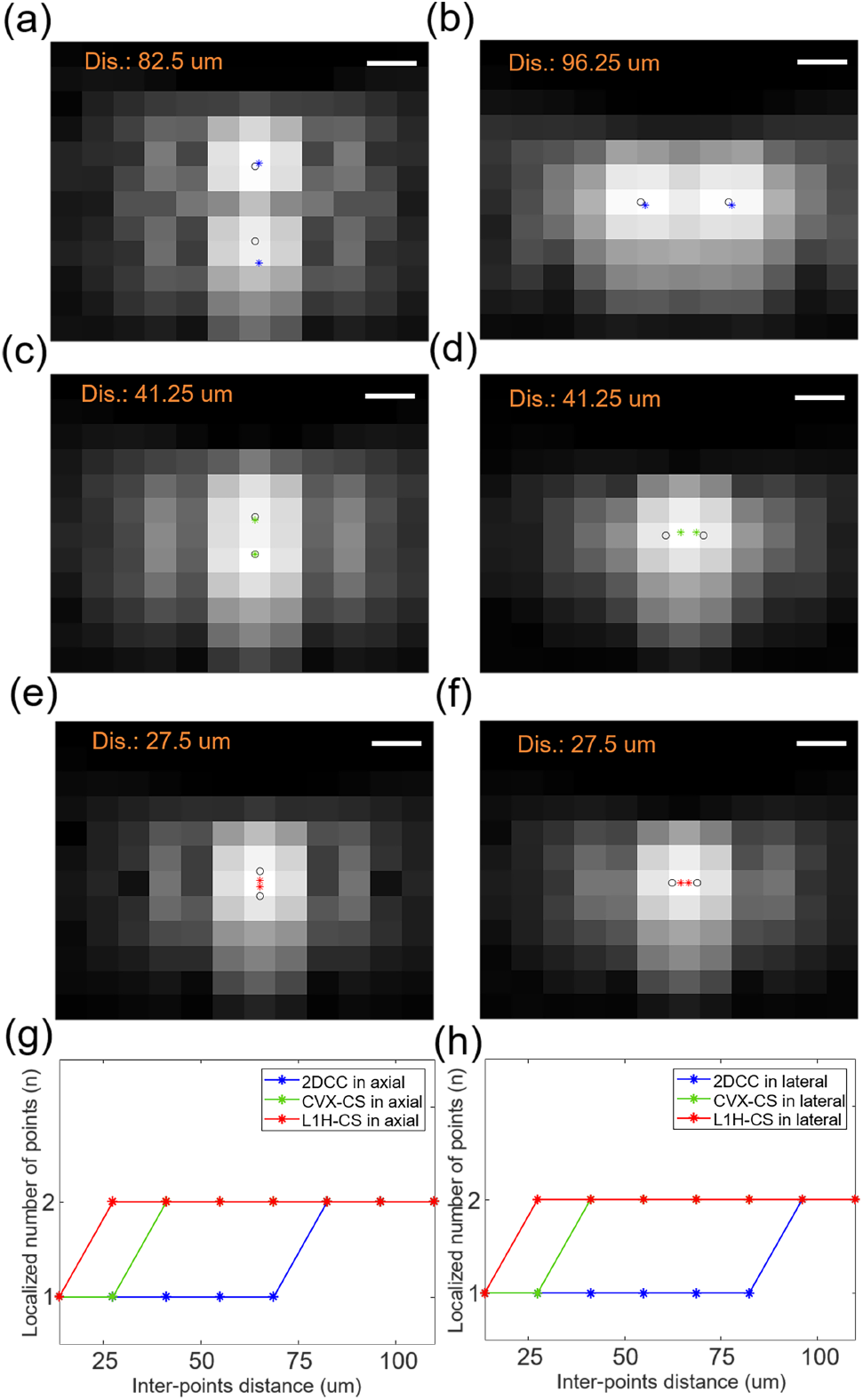
Analysis of the minimum inter-points distance between the two PTs by(a-b) 2DCC, (c-d) CVX-CS, and (e-f) L1H-CS in the axial and the lateral directions. The localization results are represented by blue, green, and red asterisks overlaid with B-mode images, while black circles show simulated locations of two PTs. Localized number of points depending on the inter-points distance and localization techniques in (g) axial and (h) lateral directions. Ideal localization number of points is 2. Scale bars indicate 50 μm.

### B. Evaluation of the localization techniques using randomly distributed point targets in simulation

We compared the performance of the localization techniques using randomly distributed PTs depending on point density. Figs. 3(a-c) show the representative localization results by 2DCC, CVX-CS, and L1H-CS overlaid with B-mode images of randomly distributed PTs at a density of 55.62 PTs/mm^2^. Blue, green, and red asterisks indicate localized points by 2DCC, CVX-CS, and L1H-CS, respectively. Black circles indicate the location of simulated points. For randomly distributed PTs at a density of 55.62 PTs/mm^2^, the average densities with the standard deviation of localized points using 2DCC, CVX-CS, and L1H-CS algorithms were measured as 21.93 ± 1.97, 46.4 ± 2.85, and 50.74 ± 0.62 PTs/mm^2^, respectively. As shown in the insets of Figs. 3(a-c), overlapped points were localized using CS algorithm, but 2DCC could not localize them. Figs. 3(d-e) show localized PTs density and localization error by 2DCC, CVX-CS, and L1H-CS. The black dashed line in Fig. 3(d) indicates the ideal localization of PTs. 2DCC algorithm departs from the black dashed line significantly when the density of simulated points reaches 15.89 PTs/mm^2^ (blue line in Fig. 3(d)). CS-based algorithms followed the black dashed line closely until the target density increased to 55.62 PTs/mm^2^ as indicated by the black arrow in Fig. 3(d). Especially, L1H-CS could localize over 90% of simulated points until the PTs density of 55.62 PTs/mm^2^. The maximum localization of 2DCC and CS algorithms are approximately 39.73 and 145.7 PTs/mm^2^ [Fig. 3(d)]. Until the target density reaches 55.62 PTs/mm^2^, localization error using CVX-CS, and L1H-CS algorithms are below 8.17 ± 0.26 and 7.38 ± 0.84 μm, while 2DCC shows the localization error approximately 12.35 ± 0.93 μm. Fig 3(f) shows computational time to localize simulated PTs. The average computational times with the standard deviation to localize points at the density of 55.62 PTs/mm^2^ using 2DCC, CVX-CS, and L1H-CS were measured to be 0.11 ± 0.01 sec, 14.75 ± 2.58 sec, and 0.76 ± 0.06 sec, respectively. L1H-CS algorithm is ~19.4× faster than the CVX-CS algorithm.

**Fig. 3.**
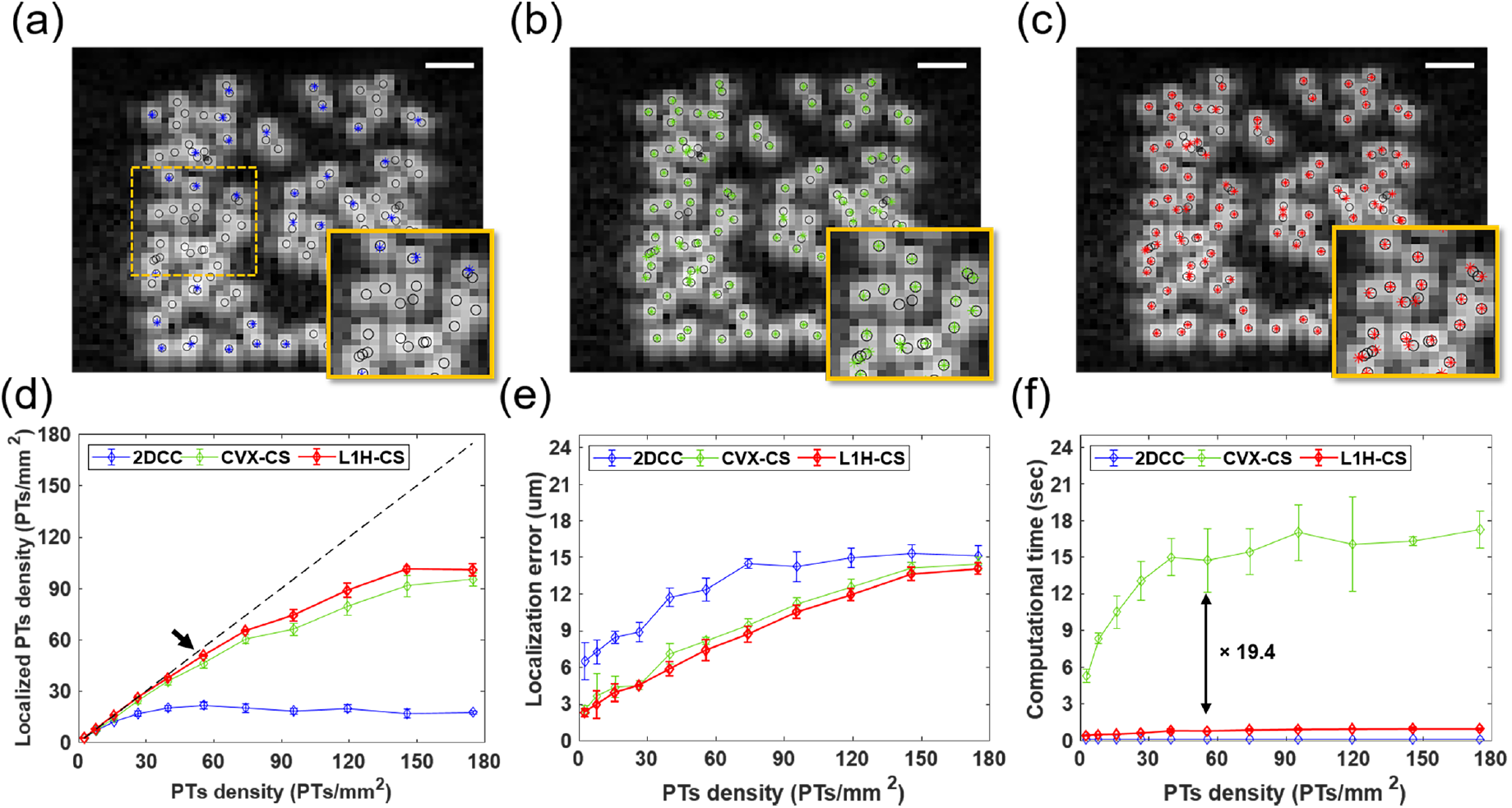
Evaluation of localization technique using randomly distributed PTs (n = 5 for each density). Localization results of PTs with a density of 55.62 PTs/mm^2^ are shown in (a) 2DCC, (b) CVX-CS, and (c) L1H-CS algorithms. Localized points by 2DCC, CVX-CS, and L1H-CS are blue, green, and red asterisks overlaid with B-mode images, while black circles show the positions of simulated PTs. The overlapped points were localized by CS algorithm, but 2DCC could not localize them as shown in the insets images of (a-c). (d) Localization efficiency, (e) error, and (f) computational time depending on PTs density from 2.65 to 174.8 PTs/mm^2^ describe that CVX-CS and L1H-CS algorithms can efficiently localize simulated PTs with 90% yield up to the density of 55.62 PTs/mm^2^. While 2DCC can localize PTs with 90% yield up to the PT density of 15.89 PTs/mm^2^. The Black dashed line indicates ideal match between simulated points and localized points by each algorithm. Image size is 62×62 pixels and scale bars indicate 250 μm.

### C. Evaluation of post-processing techniques using images of in vivo mouse tumor model

After evaluating the localization technique using simulated PTs, we first examined the effects of combinations of post-processing techniques including SVD, noise equalization, and NLM filtering to improve CTR and SNR of MBs signals. Figs. 4(a-d) show images before and after applying SVD, noise equalization, and NLM filtering to the data set acquired by ultrasound imaging of in vivo mouse tumor model. To calculate CTR, MBs and tissue signals were collected from the B-mode, SVD, SVD + noise equalization, and SVD + noise equalization + NLM filtered images. ROIs were selected at the same depth indicated as the orange boxes shown in Fig. 4(b) for all images. CTR of B-mode, SVD, SVD + noise equalization, and SVD + noise equalization + NLM filtering were measured to be - 0.23, 12.84, 13.83, and 15.1 dB, respectively, as shown in Fig. 4(e). We measured SNR of SVD, SVD + noise equalization, and SVD + noise equalization + NLM filtering along the scan line indicated by the orange dotted line in Fig. 4(b). Fig. 4(f) shows the intensity profiles along the same scan line shown in Fig. 4(b) images after each step as shown in Figs. 4(b-d). SNR improves from 21.02 to 27.35 dB as noise equalization and NLM filtering were applied as shown in Fig. 4(g).

**Fig. 4.**
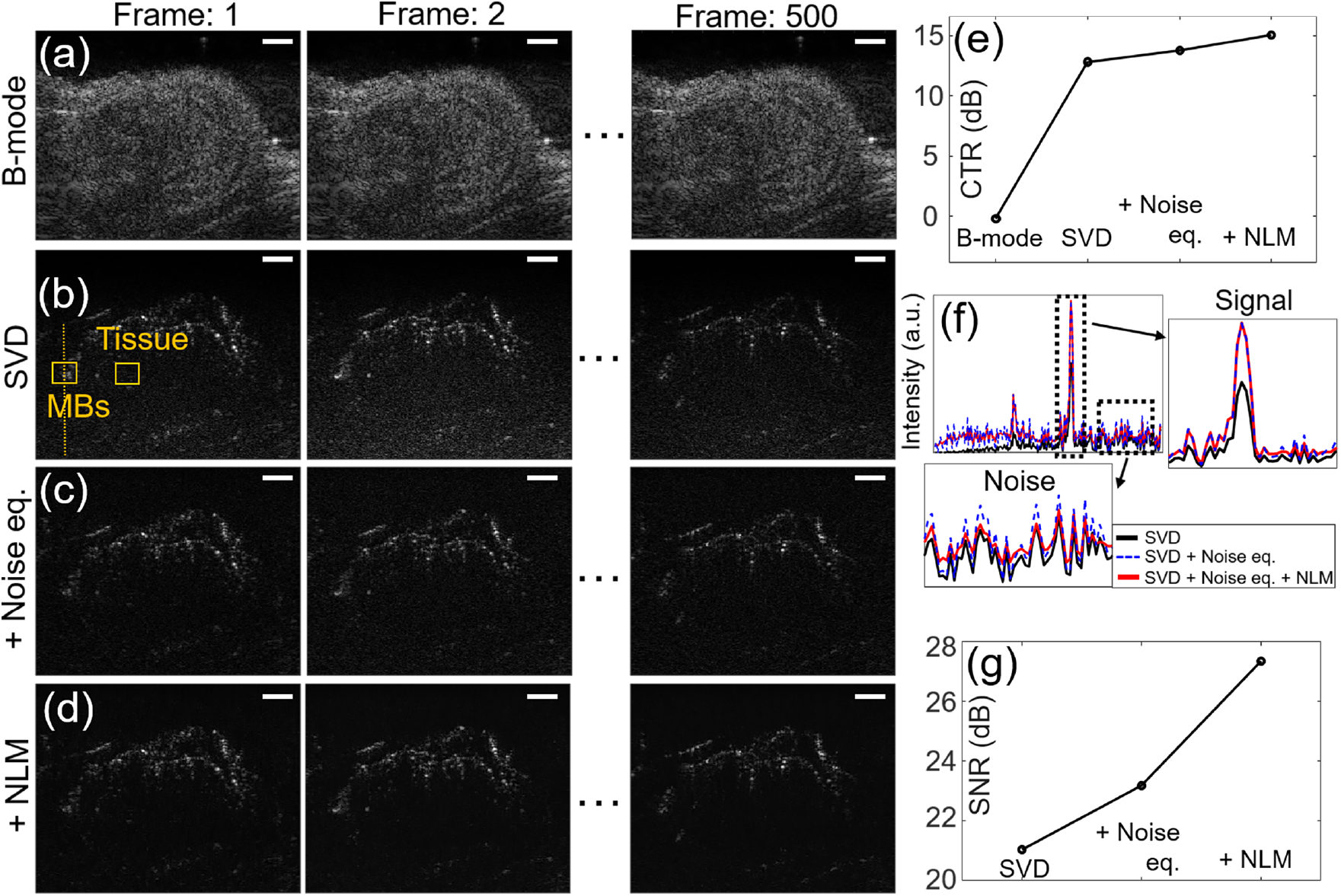
Evaluation of combination of post-processing techniques including SVD, noise equalization, and NLM filtering to the data sets acquired by ultrasound imaging of mouse tumor model after MBs injection. (a) B-mode images of frames 1 to 500 are shown. Denoised images after applying post-processing techniques including (b) SVD, (c) Noise equalization, (d) NLM filtering are presented. (e) Comparison of CTR using the acquired signals at the ROIs indicated by the orange boxes in the first frame of (a-d). (f) Intensity profiles along the scan line indicated by orange dotted line in (b-d) shows signal increase without increasing noise level. (g) SNR of images are compared. White scale bars indicate 1 mm.

Using denoised data sets, we reconstructed PWD and SRUS images using 2DCC, CVX-CS, and L1H-CS algorithms as shown in Figs. 5(a-d). As indicated by the white arrows in Fig. 5(a), smaller vessels were visualized efficiently by SRUS imaging [Fig. 5(a-d)]. We further examined the FWHM of vessels along the orange dotted line. As the black dotted line indicates in Fig. 5(e), FWHM of PWD, 2DCC, CVX-CS, and L1H-CS were measured to be 141, 82, 86, and 83 μm, respectively. The total computational time for the reconstruction of an SRUS image using the 500 frames with the image size of 252 × 252 pixels was measured to be approximately 4 min (2DCC), 4 hours (CVX-CS), and 20 min (L1H-CS), respectively [Fig. 5(f)].

**Fig. 5.**
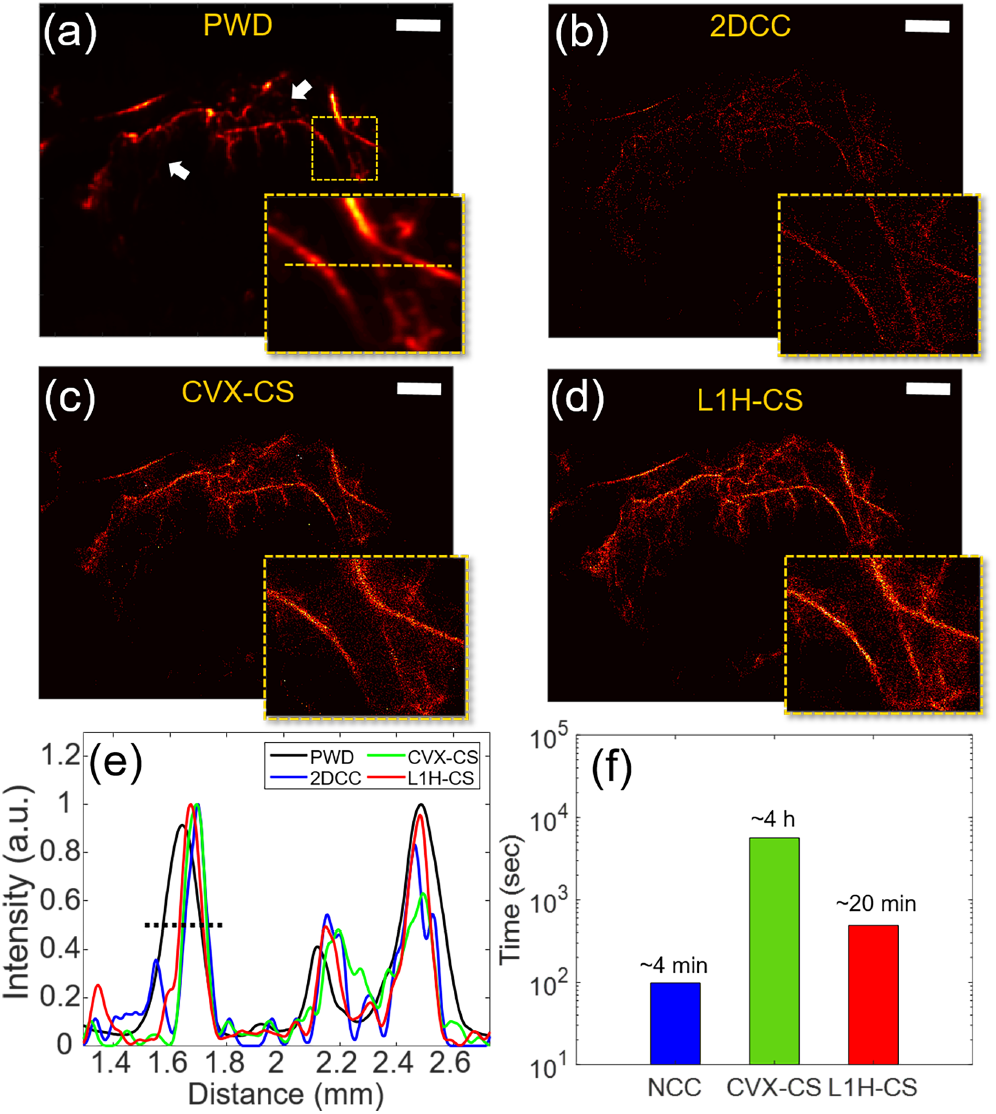
SRUS images of mouse tumor using (a) PWD, (b) 2DCC, (c) CVX-CS, and (d) L1H-CS algorithms. (e) Intensity profiles along the orange lines indicated in (a). (f) Total computational time for the image size of 252 × 252 pixels with 500 frames. White scale bars indicate 1 mm.

We further examined the effect of the number of frames by measuring the VD in the tumor. The SRUS image reconstructed by the L1H-CS algorithm using the 500 frames was overlaid with the B-mode image as shown in Fig. 6(a). The tumor area was indicated by the white dotted circles shown in Fig. 6(a). The maximum VD inside the tumor area of the SRUS image reconstructed by L1H-CS [Fig. 6(a)] with 500 frames were measured to be 13.76%. We also measure the maximum VD within the same region of images reconstructed by 2DCC and CVX-CS, which are 4.08% and 9.82%, respectively as shown in Fig. 6(b). To calculate the computational cost to acquire the similar quality of vessel images between three algorithms, we determined the number of frames to be used at VD of 4.08% as shown in Fig. 6(b). When 500 frames, 125 frames, and 75 frames were used for 2DCC, CVX-CS, and L1H-CS as shown in Figs. 6(c-e), VD of 4.08% were achieved. Computational times were measured to be approximately 4 min (2DCC), 59 min (CVX-CS), and 3 min (L1H-CS), respectively, as shown in Fig. 6(f).

**Fig. 6.**
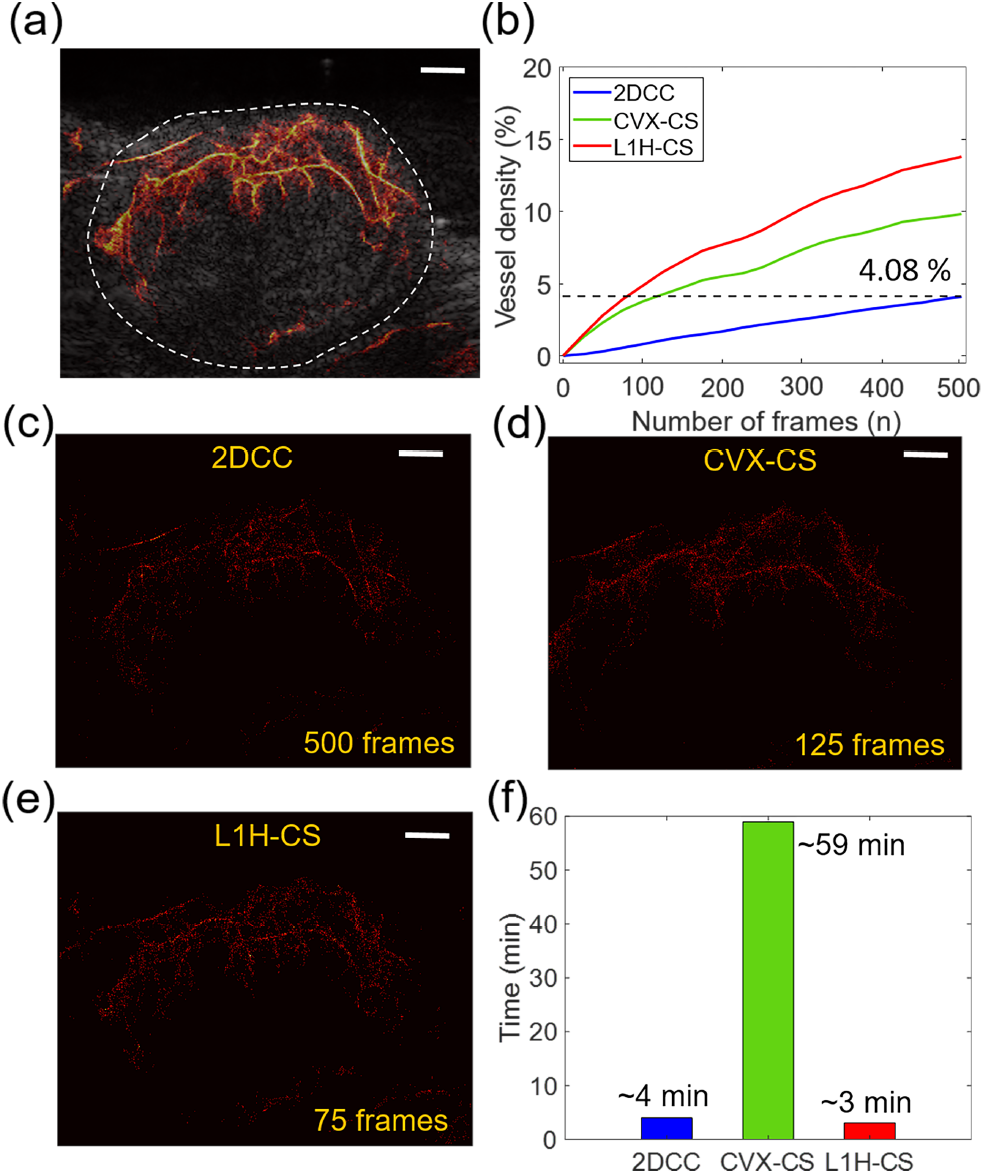
Comparison of reconstructed SRUS images of mouse tumor depending on the number of frames. (a) A SRUS image reconstructed by L1H-CS with 500 frames overlaid with B-mode image describes the vasculature inside tumor. (b) VD in the tumor area of SRUS reconstructed by three algorithms were measured depending on the number of frames. (c) SRUS images reconstructed by (c) 2DCC (500 frames), (d) CVX-CS (125 frames), and (e) L1H-CS (75 frames) shows the same VD of 4.08%. (f). Computational time to reconstruct SRUS using 500 frames for 2DCC, 125 frames for CVX-CS, and 75 frames for L1H-CS to reach the same VD. White scale bars indicate 1 mm.

### D. In vivo mouse and zebrafish super-resolution ultrasound imaging using ℓ_1_-homotopy based compressed sensing algorithm

Figs. 7(a-d) represent reconstructed SRUS image overlaid with B-mode images of mouse tumor and kidney and zebrafish dorsal trunk and brain, respectively. We measured the vessel size and distance of adjacent vessels at the ROI along the yellow lines in Figs 7(a-d). In SRUS images of mouse tumor, the vessel sizes were measured to be 25.68 μm in profile 1, and the distances between adjacent vessels were measured to be 109.9 and 109.01 μm in profiles 2 and 3, respectively [Fig. 7(e)]. The vessel sizes were measured as 32.2, and 22.2 μm in profiles 1, and 2, and the distance of adjacent vessels was estimated as 149.9 in profile 3 of the SRUS image of mouse kidney [Fig. 7(f)]. Besides, the vessel sizes were measured as 55.7, and 85.85 μm in profile 1, 64.8 μm in profile 2, and 61.8 μm in profile 3, respectively, in the SRUS image of zebrafish dorsal trunk [Fig. 7(g)]. The vessels size of 63.23 and 64.2 μm in profiles 1, and 2 were detected in the SRUS image of zebrafish brain [Fig. 7(h)]. The smallest vessel size is 22.2 μm by SRUS imaging. Considering the half wavelength of 25 MHz ultrasound as 32 μm, the diffraction limit was overcome by L1H-CS based SRUS imaging.

**Fig. 7.**
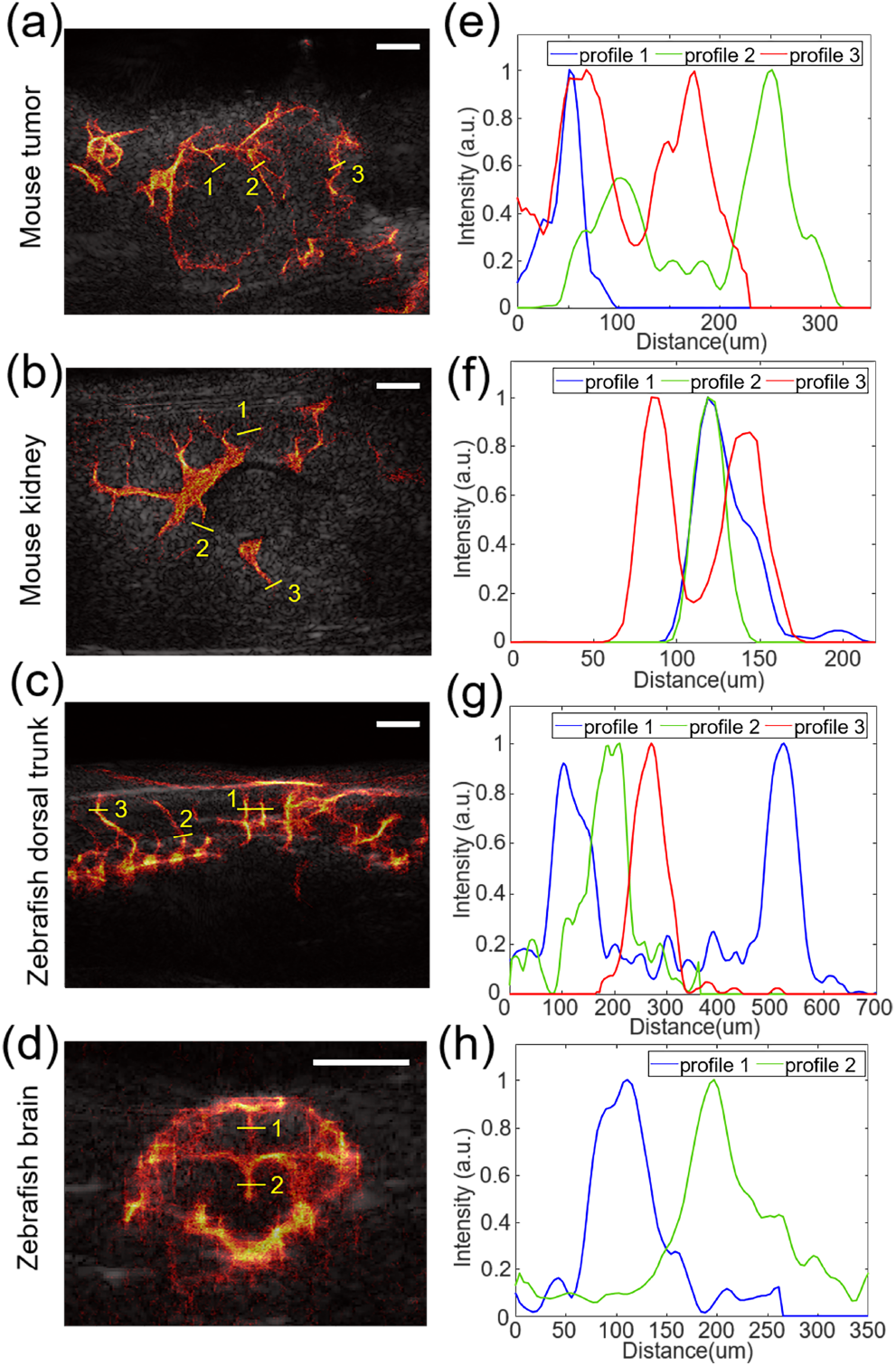
Reconstruction of SRUS images overlaid with B-mode images of mouse (a) tumor and (b) kidney and zebrafish (c) dorsal trunk and (d) brain. (e-f) Intensity profiles along the yellow lines indicated in (a-d). The smallest vessel size was measured as 22.2 μm in mouse kidney as shown in profile 2 of (f). Half wavelength of 25 MHz is 32 μm. White scale bars indicate 1 mm.

## IV. Discussions

Based on the results demonstrated in this study, L1H-CS can be used to decrease imaging acquisition time to acquire a similar quality of SRUS images [Fig. 6] or improve the image quality of SRUS with significantly improved computational cost [Fig. 5]. L1H-CS only requires 75 frames to reconstruct the similar quality, validated by VD, projected acquisition time is only 15% of that of typical 2DCC method. To acquire high quality SRUS images, validated by VD, 500 frames are used with 5 times higher computational cost. However, the computational cost of L1H-CS was decreased by 5 times compared to CVX-CS algorithm. This exhibits the potential to be used in clinics.

While L1H-CS improves acquisition time and image quality, L1H-CS also overcomes diffraction-limit as other SRUS imaging methods reported in other groups work [12, 39–41]. We were able to detect 22 μm vessels using L1H-CS in mouse kidney imaging [Fig. 7].

The localization results of the two-PTs and randomly distributed-PTs demonstrated that CS-based algorithms improved localization efficiency compared to 2DCC algorithm. L1H-CS algorithm exhibited slightly better performance than the CVX-CS algorithm to localize the PTs. In particular, CVX-CS localized two-PTs when the distance between two points approaches 27.5 μm, on the other hand, L1H-CS algorithm localized two points when their distance approaches 13.75 μm [Fig. 2]. Randomly distributed PTs with a density of 55.62 PTs/mm^2^ were localized to be 50.74 ± 0.62 PTs/mm^2^ by L1H-CS, which demonstrated 10% localization accuracy compared to CVX-CS. One possible explanation of improved localization efficiency of L1H-CS compared to CVX-CS is that L1H-CS algorithm searches optimized regularization parameter *ϵ*, while CVX-CS fixed the constant *ε* as 2.3 [Eqs. (2) and (4)] [21–23].Besides, L1H-CS required a much lower computing cost than the CVX-CS algorithm [Fig. 3(f) and Fig. 5(f)]. This is because the computational complexity of L1H-CS is no longer affected by the inverse matrix, which is a major factor that demands the high computational cost in CVX-CS algorithm [28].

Commonly used 2DCC for SRUS imaging technique could not localize if PTs or MBs are closely located as shown in Fig. 2 and Fig. 3. To track individual MBs efficiently in vessels, lower MBs concentration was usually used resulting in long acquisition time [7, 8]. L1H-CS localization technique proposed here can detect densely populated PTs or MBs efficiently as demonstrated in Fig. 2 and Fig. 3. Thus, we performed the SRUS imaging of mouse and zebrafish by injecting the high-density MBs (0.75 ‒ 2.0 × 109 MBs/mL) than the standard density of MBs (1.5 – 5.6 × 108 MBs/mL). In the in vivo experiment, VD in tumors was measured as 4.08% based on microvessel image reconstructed by 2DCC, while L1H-CS could reconstruct microvessel of 13.76% in tumor regions using 500 frames [Fig. 6(b)]. These results suggest that the L1H-CS algorithm allows SRUS imaging with the injection of high-density MBs, suggesting that the acquisition time of ultrasound image frames can be reduced, resulting in a faster imaging session would be implemented.

PWD image could not represent microvessels indicated by white arrows as shown in Fig. 5(a), while SRUS images showed the microvessel at the same regions [Figs. 5(b-d)]. This is because of the lack of the sensitivity of the PWD imaging technique when MBs flows in a very tiny vessel, resulting in showing the large gap of the contrast between the large vessels and tiny vessels. Thus, the tiny vessels might not be visualized in PWD imaging. Proposed L1H-CS may be used to investigate tumor micro-vascularization as the tumor progresses.

## V. CONCLUSIONS

In conclusion, we described the L1H-CS based SRUS imaging technique that could reduce computational time and acquisition time or significantly improve SRUS image quality with some expenses of post-processing cost. In the simulation, we confirmed that a minimum distinguishable distance of PTs was 13.75 μm, which is a sub-diffraction limit and efficient localization of PTs compared to a conventional localization technique. By in vivo mouse imaging, the minimum detectable blood vessel size in mouse kidney was measured to be 22 μm, which confirms the sub-diffraction limit capability of L1H-CS. Based on results both in simulation and in vivo experiments, the proposed algorithm significantly improves SRUS image quality and data acquisition time.

